# Genomewide significant regions in 43 Utah high-risk families implicate multiple genes involved in risk for completed suicide

**DOI:** 10.1101/195644

**Authors:** Hilary Coon, Todd M. Darlington, Emily DiBlasi, W. Brandon Callor, Elliott Ferris, Alison Fraser, Zhe Yu, Nancy William, Sujan C. Das, Sheila E. Crowell, Danli Chen, John S. Anderson, Michael Klein, Leslie Jerominski, Dale Cannon, Andrey Shabalin, Anna Docherty, Megan Williams, Ken R. Smith, Brooks Keeshin, Amanda V. Bakian, Erik Christensen, Qingqin S. Li, Nicola J. Camp, Douglas Gray

## Abstract

Suicide is the 10th leading cause of death in the US. While environment has undeniable impact, evidence suggests genetic factors play a significant role in completed suicide. We linked a resource of >4,500 DNA samples from completed suicides obtained from the Utah Medical Examiner to genealogical records and medical records data available on over 8 million individuals. This linking has resulted in the identification of high-risk extended families (7-9 generations) with significant familial risk of completed suicide. Familial aggregation across distant relatives minimizes effects of shared environment, provides more genetically homogeneous risk groups, and magnifies genetic risks through familial repetition. We analyzed Illumina PsychArray genotypes from suicide cases in 43 high-risk families, identifying 30 distinct shared genomic segments with genome-wide evidence (p=2.02E-07 to 1.30E-18) of segregation with completed suicide. The 207 genes implicated by the shared regions provide a focused set of genes for further study; 18 have been previously associated with suicide risk. While PsychArray variants do not represent exhaustive variation within the 207 genes, we investigated these for specific segregation within the high-risk families, and for association of variants with predicted functional impact in ~1300 additional Utah suicides unrelated to the discovery families. None of the limited PsychArray variants explained the high-risk family segregation; sequencing of these regions will be needed to discover segregating risk variants, which may be rarer or regulatory. However, additional association tests yielded four significant PsychArray variants (*SP110*, rs181058279; *AGBL2*, rs76215382; *SUCLA2*, rs121908538; *APH1B*, rs745918508), raising the likelihood that these genes confer risk of completed suicide.

## Introduction

Suicide is the 10th leading cause of death in the United States; over 44,000 individuals die by suicide in the US every year.^1^ While environmental variables have undeniable impact, evidence suggests that genetic factors play a role in completed suicide, with heritability of close to 50%.^2,3^ Recent growth in the number of suicide genetic studies has resulted in promising findings from candidate gene and genome-wide association studies,^4^ though many remain to be replicated. Replication is hampered by sample differences across studies, including differences in demographics and primary diagnoses of study samples, as many studies of suicide risk have been conducted within cases ascertained for specific psychiatric disorders.^4^ In addition, most studies of suicide have focused on suicidal ideation and behaviors; these phenotypes are much more common than completed suicide, allowing for ascertainment of sufficiently powered samples, but suicidal behaviors can be difficult to quantify, and represent individuals with a range of risk for later suicide. In addition, evidence suggests important differences in the etiology of suicidal behaviors versus the less ambiguous but much rarer outcome of completed suicide.^5^

We implemented a unique study design to investigate genetic risk for suicide through the collection of DNA samples on >4,500 consecutive individuals who died by suicide in the state of Utah, providing an unparalleled population-based genetic resource. This sample results from a long-term collaboration with the Utah State Office of the Medical Examiner. The records from these cases have been linked to the Utah Population Database (UPDB, https://healthcare.utah.edu/huntsmancancerinstitute/research/updb), a comprehensive database including multi-generational genealogies, as well as death certificates, demographic data, and current medical information on over 8 million individuals. Through this linking we have identified very large families (7-9 generations) with significantly elevated suicide risk. Familial aggregation across distant relatives in these families minimizes the impact of shared environment on risk. High-risk families also provide more genetically homogeneous risk groups, increasing statistical power to detect familial variants associated with disease risk. The Utah extended family study design has already shown success in the study of other complex genetic diseases of extended families of similar size (e.g., colon cancer,^6^ breast cancer,^7^ and cardiac arrhythmia^8^).

This study reflects an analysis of 43 very large Utah families at significantly elevated risk for completed suicide. The focus on completed suicide, statistically concentrated in these high-risk families, optimizes power to reveal regions of the genome likely to contain risk variants. Our design investigates genetic risk using the suicide cases in the extended families regardless of co-occurring psychopathology, and continues with follow-up studies from our population-wide ascertainment of all suicide deaths in Utah, again without regard to co-occurring psychopathology. We recognize that psychiatric diagnoses are critically important in suicide risk;^9,10^ it is likely that findings from our study are related to these associated risks. However, because of the familial aspect of our design, it is possible that our results may reveal risk variants that cross-cuts specific psychiatric diagnoses.^11–14^

Genetic studies of psychiatric disease have revealed associations with multiple rare and common risk variants with reduced penetrance, which may interact in complex ways with each other, with background genetics, and with environmental risks.^15^ Based on results to date, we expect that suicide will follow this complex genetic architecture. In this study, the familial analyses use a new statistical method (Shared Genomic Segments, SGS^17^) that is well-powered to identify rare genetic variants in large families, evidence that can then be used to prioritize searches for additional variants contributing to risk in other case samples. This design is complementary to the Genome-Wide Association Study (GWAS) approach in large case-control samples, which can also produce statistical evidence for risk genes to be followed up in independent case samples.

Using genome-wide single nucleotide polymorphism (SNP) variants matched to the same variants in publicly-available population control data, we identified regions of the genome that segregate in suicide cases within high-risk families. These statistically significant regions provide compelling genes as targets for follow up. While such follow-up studies would ideally use comprehensive sequence data, the SNP array platform used in this study contains putatively functional variants of high interest to psychiatric and medical disorders, both of which may share overlapping suicide risk. Functional content of the PsychArray was investigated first within familial cases responsible for the significant regions, and then within ~1300 additional Utah suicide cases unrelated to the original extended families analyzed.

This study adds to the growing knowledge of genetic risk for suicide. First, we have identified genes in regions of significant familial segregation in large high-risk families, providing replication for previously reported genes of high interest, and identifying target genes for additional follow up. Second, we have identified novel risk variants using a large follow-up association analysis of PsychArray variants with predicted functional impact using a population-matched resource of suicide cases.

## Materials & Methods

### Sample

This project is possible because of a collaboration with the Utah State Office of the Medical Examiner (OME) which has spanned two decades. With Institutional Review Board (IRB) permissions from the University of Utah, the Utah Department of Health and Utah Intermountain Healthcare, we have collected de-identified DNA samples from consecutive suicides since 1997. The collection numbers 4,585 (3,632 males and 953 females). DNA was extracted from blood using the Qiagen Autopure LS automated DNA extractor (www.qiagen.com). Identifying information from cases with DNA was linked to data within the UPDB’s secure computer servers. All identifying data was then stripped before providing data to the research team; suicide cases and family structure data are referenced by anonymous IDs. DNA for this research project is shared with the NIMH Repository and Genomics Resource, project number 315 (2,880 samples are now at the repository; additional samples are being sent on an ongoing basis).

### Determination of familial risk, selection of families/cases

Genealogical data in the UPDB was used to construct family trees and identify those families at high risk for suicide. Beyond the suicide cases with DNA, the UPDB contains records of all known suicides from Utah death certificates dating from 1904 (N=14,288). All 14,288 cases were used to estimate familial risk of suicide. To determine the extended families at highest risk, we used the Familial Standardized Incidence Ratio (FSIR) statistic,^16^ calculated by comparing the incidence of suicide in each extended family to its expected incidence determined by the statewide distribution for suicide stratified by sex and age. We identified 241 high-risk families containing significant excess of suicides (p<0.05) and at least three suicides with DNA. We selected 43 of these families for analysis (Table 1) based on significance of the FSIR risk statistic, number of cases with DNA, and overall count of meioses between these cases (see Table 1). The 43 families included 2.04-4.41 times the expected number of suicide cases as reflected in the FSIR statistic (range p=0.003 to 1E-12, average p=0.0007). The average number of cases per family with genotyping was 6.2 (range 3-13), and the average number of meioses between analyzed cases was 29.6 (range 15-70; see Figure 1 for an example of how meioses are counted in a family of moderate size from our resource). Family-specific significance thresholds for genomic sharing (see analysis section below) depend upon family size, structure, dispersion of cases, and number of cases analyzed. Permission for use of family structure data was granted by the Resource for Genetic and Epidemiologic Research (RGE, https://rge.utah.edu), the oversight committee for use of UPDB data.

**Table 1.** Characteristics of 43 extended families at high risk for suicide.

| Family | FSIR | FSIR P-value | Total N Obs. Cases | Total N Exp. Cases | N Cases in Study | N Meioses Between Analyzed Cases | SGS Threshold: Significant | SGS Threshold: Suggestive | ≥ 1 Significant Region | ≥ 1 Suggestive Region |
| --- | --- | --- | --- | --- | --- | --- | --- | --- | --- | --- |
| 709 | 2.69 | <0.0001 | 27 | 10.04 | 8 | 35 | 6.36E-08 | 6.41E-07 | YES | YES |
| 1881 | 3.12 | <0.0001 | 21 | 9.31 | 4 | 17 | 2.03E-06 | 3.00E-05 |  |  |
| 2082 | 3.99 | <0.0001 | 19 | 4.77 | 3 | 15 | 3.93E-06 | 9.78E-05 |  |  |
| 7785 | 2.78 | <0.0001 | 28 | 10.08 | 6 | 26 | 1.45E-07 | 1.82E-06 |  | YES |
| 8556 | 2.85 | <0.0001 | 41 | 14.39 | 7 | 37 | 8.99E-08 | 1.01E-06 |  | YES |
| 11593 | 2.31 | 0.0002 | 25 | 10.82 | 6 | 29 | 1.28E-07 | 1.70E-06 |  | YES |
| 12291 | 2.43 | 0.0001 | 23 | 9.48 | 6 | 29 | 1.59E-07 | 1.96E-06 |  | YES |
| 27251 | 2.43 | 0.0001 | 36 | 16.93 | 12 | 52 | 1.16E-09 | 1.41E-08 |  | YES |
| 36667 | 3.61 | 0.0039 | 7 | 1.61 | 4 | 17 | 7.99E-07 | 1.56E-05 |  | YES |
| 37661 | 2.06 | 0.0031 | 19 | 9.21 | 5 | 20 | 1.79E-06 | 1.91E-05 |  | YES |
| 40780 | 3.61 | 0.0038 | 7 | 1.94 | 4 | 19 | 1.59E-06 | 2.57E-05 |  | YES |
| 41469 | 2.29 | 0.0013 | 18 | 7.86 | 4 | 20 | 4.24E-06 | 5.86E-05 |  | YES |
| 43035 | 2.46 | <0.0001 | 36 | 14.63 | 7 | 31 | 1.09E-07 | 1.15E-06 |  | YES |
| 43580 | 2.45 | 0.0004 | 19 | 7.75 | 6 | 29 | 2.93E-07 | 3.08E-06 |  | YES |
| 46547 | 2.48 | <0.0001 | 29 | 11.71 | 6 | 28 | 2.75E-07 | 2.92E-06 |  | YES |
| 60205 | 2.26 | 0.0007 | 21 | 9.31 | 5 | 26 | 5.07E-07 | 6.52E-06 | YES |  |
| 66494 | 2.45 | 0.0001 | 24 | 9.8 | 7 | 34 | 5.45E-08 | 6.47E-07 | YES | YES |
| 68939 | 2.21 | 0.0007 | 22 | 9.97 | 8 | 41 | 3.66E-08 | 3.95E-07 |  | YES |
| 91500 | 3.19 | 0.0003 | 13 | 4.07 | 4 | 16 | 1.24E-06 | 2.18E-05 |  |  |
| 129334 | 2.75 | <0.0001 | 25 | 9.09 | 7 | 36 | 6.92E-08 | 8.32E-07 |  | YES |
| 148039 | 3.65 | 0.0002 | 12 | 3.29 | 4 | 15 | 5.14E-06 | 5.73E-05 |  |  |
| 176860 | 2.71 | <0.0001 | 43 | 15.88 | 9 | 39 | 3.28E-08 | 3.08E-07 |  | YES |
| 185855 | 2.96 | 0.0003 | 15 | 5.06 | 6 | 25 | 2.46E-07 | 2.24E-06 |  | YES |
| 209487 | 2.82 | <0.0001 | 26 | 9.23 | 7 | 32 | 1.19E-07 | 1.25E-06 | YES | YES |
| 233769 | 2.5 | <0.0001 | 33 | 13.19 | 11 | 50 | 1.58E-08 | 1.40E-07 | YES | YES |
| 265545 | 4.05 | 0.001 | 8 | 1.98 | 5 | 17 | 2.74E-06 | 3.82E-05 |  |  |
| 540295 | 2.59 | <0.0001 | 43 | 16.59 | 5 | 31 | 1.73E-07 | 3.10E-06 |  | YES |
| 540775 | 2.46 | 0.003 | 13 | 5.28 | 7 | 34 | 5.88E-08 | 7.45E-07 | YES | YES |
| 544252 | 3.21 | 0.0025 | 9 | 2.8 | 4 | 19 | 7.48E-07 | 1.43E-05 |  | YES |
| 553615 | 2.04 | <0.0001 | 81 | 39.7 | 13 | 69 | 2.69E-10 | 3.74E-09 | YES | YES |
| 554151 | 2.38 | 0.003 | 14 | 5.89 | 7 | 32 | 9.98E-08 | 1.05E-06 |  |  |
| 587072 | 2.39 | 0.003 | 14 | 5.86 | 5 | 25 | 1.39E-06 | 1.55E-05 |  | YES |
| 590241 | 2.47 | 0.0004 | 19 | 7.7 | 8 | 38 | 2.86E-08 | 3.16E-07 |  | YES |
| 595955 | 2.48 | <0.0001 | 39 | 15.73 | 7 | 39 | 5.35E-08 | 6.62E-07 |  | YES |
| 601627 | 2.86 | <0.0001 | 69 | 24.14 | 12 | 70 | 7.06E-10 | 9.14E-09 | YES | YES |
| 603481 | 2.64 | 0.0003 | 18 | 6.83 | 9 | 37 | 5.09E-08 | 5.35E-07 | YES | YES |
| 622459 | 2.54 | 0.0001 | 22 | 8.68 | 4 | 18 | 1.75E-06 | 2.60E-05 |  | YES |
| 755858 | 2.28 | 0.001 | 19 | 8.33 | 5 | 24 | 3.77E-07 | 5.48E-06 |  |  |
| 756794 | 2.7 | 0.0002 | 18 | 6.68 | 5 | 24 | 3.94E-07 | 5.67E-06 |  | YES |
| 791533 | 3.67 | 0.001 | 9 | 2.45 | 3 | 17 | 3.09E-06 | 8.38E-05 |  | YES |
| 807334 | 2.5 | 0.0001 | 24 | 9.58 | 6 | 30 | 3.25E-07 | 3.57E-06 | YES | YES |
| 923763 | 4.41 | <0.0001 | 14 | 3.18 | 3 | 15 | 3.33E-06 | 8.86E-05 |  |  |
| 957634 | 3.7 | 0.0001 | 12 | 3.24 | 3 | 15 | 3.22E-06 | 8.40E-05 |  | YES |

**Figure 1.**
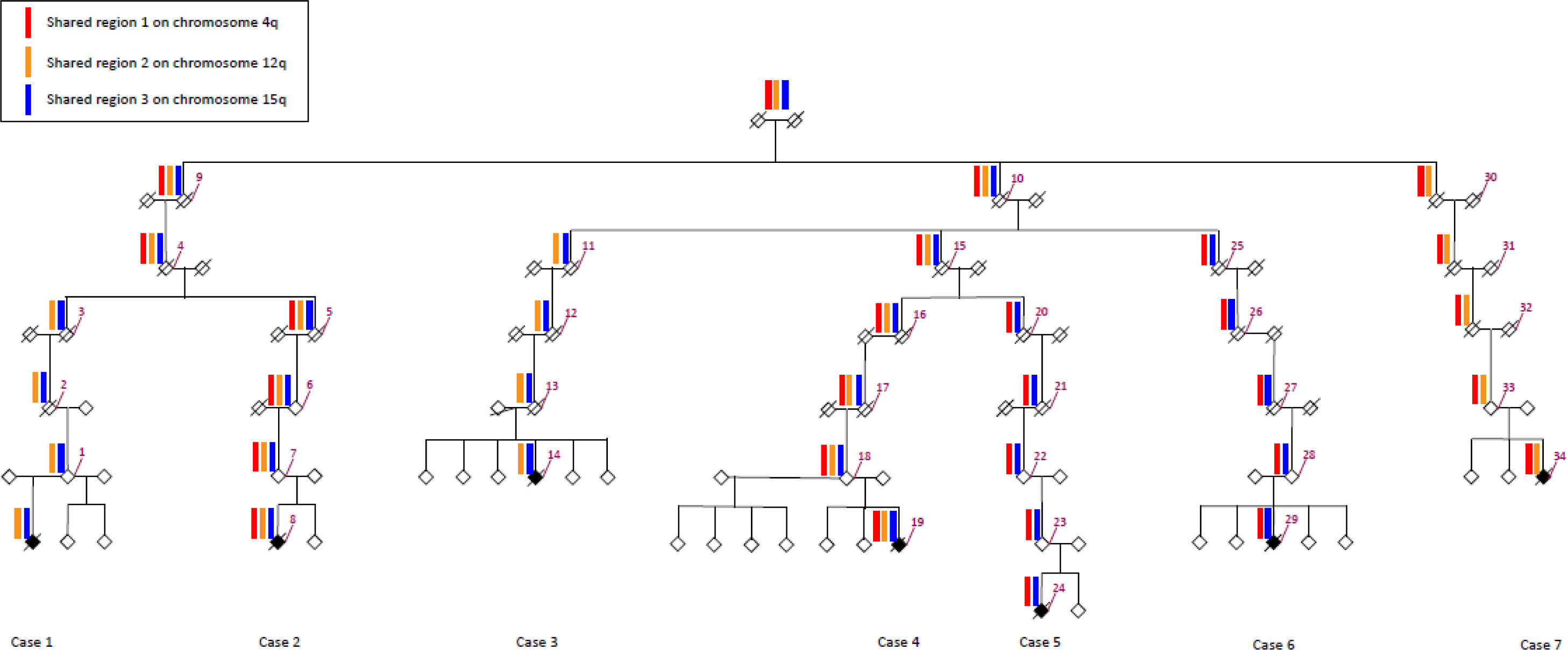
Extended structure of Family 66494 that links seven suicides (shaded in black) used for Shared Genomic Segment (SGS) analyses. Suicide cases are not as evident in upper generations because suicide status from death certificates is only available back to 1904. Note that gender is disguised and sibship order is randomized in order to protect the privacy of family members. **Family size**. There are 34 total meioses between the seven cases in this family; this counting is shown in purple on the drawing. SGS requires a total of at least 15 meioses between cases for adequate statistical power. **Shared segments**. Three genomic segments provided significant evidence of sharing between cases in this family. The pattern of segregation of each segment is shown. Cases 2, 4, 5, 6, and 7 share region 1 (red). Cases 1, 2, 3, 4, 5, and 6 share region 2 (gold). Cases 1, 2, 3, 4, and 7 share region 3 (blue). Essential segregation is shown; however, when cases *do not* share, the region can actually be lost at any meiosis above the case in the family tree. The exact point of this loss is unknown.

The 198 families not selected for analysis in this study exhibited less significant risk, and/or had too few cases with DNA, and/or had insufficient distance between cases. The average number of suicide cases per family with DNA in these 198 families was 3.23 (SD=0.88, range 3 to 5). The average p-value associated with these FSIRs was 0.0028 (SD=0.0074, range 0.0458 to 1E-4).

### Diagnostic data

In addition to basic demographic and cause of death information, we had access to diagnostic data for psychiatric conditions associated with suicide using electronic medical records data through the UPDB. Codes were linked to case numeric case identifiers within the Utah Population Database; de-identified results were provided for analysis. Conditions were defined by groups of diagnostic codes aggregated according to the International Classification of Diseases (ICD) system (www.icd9data.com; see Supplemental Table S1 for the list of codes used to define diagnostic categories in this study). Importantly, because our cases are not derived from a clinical population, cases can exhibit with no co-occurring diagnoses. Missing diagnostic data can occur for many reasons, including: 1) existence of diagnostic codes other than the 359 codes in Table S1; 2) a case who did not seek medical attention for the psychiatric disorders in question due to stigma, lack of insurance, cultural barriers or other lack of access to services, age-related lack of recognition of pathology, or symptoms not perceived to require medical attention; 3) diagnostic data not contained in the UPDB, including diagnoses prior to the storage of electronic diagnoses, or diagnoses given out of state or outside the ~85% coverage of electronic medical records data available in the UPDB. We treated missing data as unknown rather than assuming the absence of pathology.

**Table 2.** SGS regions with 1) genome-wide significant evidence, or 2) overlapping evidence in more than one family meeting at least suggestive significance.^a^

| Sharing Families | Chromosome | Start | End | Region length | N sharing Cases | P-Value<sup>a</sup> |
| --- | --- | --- | --- | --- | --- | --- |
| 709, 8556 | 1p34.2 | 40,433,771 | 40,555,321 | 121,550 | 6, 5 | 3.47E-12 |
| 791533, 540775 | 1q31.1-q31.2 | 190,694,813 | 191,590,362 | 895,549 | 3, 7 | 4.63E-09 |
| 601627 | 2p16.3 | 50,902,522 | 51,820,543 | 918,021 | 6 | 1.94E-10 |
| 176860, 11593 | 2q32.2 – q32.3 | 191,029,604 | 192,020,729 | 991,125 | 5, 6 | 4.31E-12 |
| 601627 | 2q36.3 – q37.1 | 230,899,765 | 231,454,354 | 554,589 | 6 | 2.39E-10 |
| 553615 | 3p14.1 | 64,735,531 | 65,289,530 | 553,999 | 8 | 8.87E-11 |
| 129334, 11593 | 3q26.33 | 181,074,751 | 181,229,833 | 155,082 | 4, 5 | 7.94E-12 |
| 603481 | 4q26 | 117,379,825 | 118,257,841 | 878,016 | 7 | 3.08E-08 |
| 807334 | 4q28.3 | 131,561,136 | 132,902,055 | 1,340,919 | 5 | 2.02E-07 |
| 8556, 66494 | 4q35.1 – q35.2 | 187,072,383 | 187,513,585 | 441,202 | 4, 5 | 1.82E-12 |
| 553615<sup>b</sup> | 5q23.3 – q31.1 | 129,199,151 | 131,819,921 | 2,620,770 | 9 | 2.39E-10 |
| 553615, 603481, 176860<sup>c</sup> | 5q23.3 – q31.1 | 129,684,909 | 131,819,921 | 2,135,012 | 7, 7, 7 | 1.30E-18 |
| 601627 | 5q33.3 – q34 | 159,633,484 | 160,328,128 | 694,644 | 7 | 5.47E-10 |
| 553615 | 6q11.1 – q12 | 62,563,817 | 64,139,997 | 1,576,180 | 8 | 1.34E-10 |
| 60205 | 6q24.3 | 148,162,328 | 148,621,930 | 459,602 | 5 | 4.02E-07 |
| 601627 | 7p21.2 | 14,144,663 | 15,001,308 | 856,645 | 6 | 2.04E-10 |
| 957634, 595955 | 7q36.1 | 150,239,676 | 151,123,529 | 883,853 | 3, 4 | 6.44E-11 |
| 587072, 595955 | 8p23.1 | 9,157,884 | 10,032,894 | 875,010 | 4, 5 | 4.71E-11 |
| 233769 | 10p15.3 | 2,408,852 | 2,881,331 | 472,479 | 7 | 1.11E-09 |
| 11593, 8556 | 10p12.33 | 17,391,660 | 17,576,227 | 184,567 | 4, 5 | 3.11E-11 |
| 27251, 233769 | 10q21.3 | 67,735,584 | 68,057,063 | 321,479 | 7, 5 | 8.22E-15 |
| 209487 | 11p11.2 – q12.1 | 47,312,689 | 56,518,769 | 9,206,070 | 6 | 6.60E-08 |
| 540775 | 11q13.3 | 69,482,091 | 69,933,696 | 451,605 | 7 | 6.20E-09 |
| 209487, 66494 | 12q.12 | 41,899,312 | 42,298,882 | 399,570 | 5, 5 | 2.14E-12 |
| 709 | 13q12.3 | 29,886,987 | 30,492,217 | 605,320 | 7 | 1.86E-08 |
| 27251, 41469 | 13q14.2 | 48,526,833 | 49,283,795 | 756,962 | 5, 4 | 2.93E-12 |
| 590241, 601627 | 14q23.1 – q23.2 | 60,699,751 | 62,360,464 | 1,660,713 | 5, 8 | 5.91E-14 |
| 709 | 15q21.3 – q22.2 | 58,601,804 | 59,646,991 | 1,045,187 | 6 | 2.74E-08 |
| 66494 | 15q22.2 | 62,914,165 | 63,686,327 | 772,162 | 6 | 5.44E-08 |
| 27251, 233769 | 18q11.2 | 24,414,687 | 24,494,344 | 79,657 | 8, 6 | 5.22E-15 |
| 27251, 622459 | 19q13.12 | 35,836,530 | 36,136,449 | 299,919 | 8, 3 | 2.89E-12 |

<sup>a</sup> For regions shared by > 1 family, p-value was estimated using Fisher's combined probability test.<sup>20</sup>
<sup>b</sup> This region was significantly shared by 553615 on its own, but a smaller overlapping region was also shared by 553615, 603481, and 176860.
<sup>c</sup> Two cases were omitted from family 553615 to satisfy the independence requirement for computing Fisher's combined p-value.<sup>20</sup> Significance thresholds were re-computed for family 553615 eliminating these cases. Person 112304 is a descendant of both 553615 and 603471, and person 95765 is a descendant of both 553615 and 176860.

### Molecular data

The SGS analyses used variants from the Illumina Infinium PsychArray platform, version 1.0 (https://www.illumina.com/products/by-type/microarray-kits/infinium-psycharray.html) genotyped on 216 suicide cases in the 43 selected families. This PsychArray includes 265,000 common informative tag SNPs, 245,000 variants selected from exome sequencing studies of medical and psychiatric conditions, and 50,000 rare variants associated specifically with psychiatric conditions. Supplemental Figure S1 shows the use of genotype data for the study. Genotyped array content was oriented to 1000 Genomes Project data. For initial analyses of familial sharing, we included all variants contained in 1000 Genomes Project control data, omitting variants where orientation was ambiguous, and variants which were not polymorphic. Using PLINK,^17^ we also removed 17,058 variants with >5% missing calls and 176 variants that failed Hardy-Weinberg equilibrium (p<0.001). Additionally, one case from family 553615 was removed due to a low call rate (>5% missing). Our initial familial analyses used 237,415 variants from 215 completed suicide cases to reveal familial variation that defines the boundaries of the segments shared among related cases. Rare putatively functional array variants meeting QC criteria, including psychiatric and medical disease-specific variants, were used to follow up additional variants in the shared regions.

### Analysis (see Supplemental Figure S1 for a flow diagram of the study)

We began by using a new analytical method, Shared Genomic Segments (SGS)^18^ developed for analyzing large high-risk families to identify subsets of cases that share regions beyond sharing expected by chance. SGS identifies excessive lengths of consecutive SNPs with allelic sharing between relatives to infer genomic segments that are inherited. Theoretically, chance inherited genomic sharing in distant relatives is extremely improbable; thus, the method has power in large families such as those in our study.^19^ The significance of each shared segment is assessed empirically using gene-drop simulations (independent of case status) to create a null distribution of expected sharing within each family. The method assigns haplotypes to family founders according to a publicly available linkage disequilibrium map from 1000 Genomes European data, followed by simulated segregation through each specific family structure, repeated a minimum of 500,000 times. The observed sharing is compared to simulated sharing to determine significance. See Supplemental Figure S2 for a hypothetical simplified example of SGS sharing. Genome-wide significance thresholds are calculated specific to each family, as statistical power varies with family structure, number of cases, and distance between cases. Significance thresholds account for multiple testing and linkage disequilibrium, and also adjust for within-family heterogeneity by including adjustment for all possible subsets of within-family sharing among cases. Model fitting to determine theoretical genome-wide thresholds used these distributions of gene-drop results. The overwhelming majority of the genome will be null (does not contain a suicide risk variant); we acknowledge a slight conservative bias as these distributions also contain a small number of true positives.^17^ The genome-wide significant threshold corresponds to a false positive rate of 0.05 per genome per family, while the suggestive threshold corresponds to one false positive result per genome per family. In this study, we report regions with family-specific genome-wide significant evidence, and regions overlapping in more than one family where family-specific evidence was at least genome-wide suggestive. P-values for these overlapping multiple-family regions were approximated using Fisher’s combined probability test.^20^ The Shared Genomic Segment (SGS) analysis software is freely available (https://uofuhealth.utah.edu/huntsman/labs/camp/analysis-tool/shared-genomic-segment.php).

Power of SGS was previously investigated for a range of genetic models involving rare variants in extended family data,^19^ showing appropriate power for large families with at least 15 meioses between cases. For all scenarios considered in this study, genome-wide association studies would have had negligible power. Given these results, we selected only extended families with at least 15 meioses between cases (see Table 1 and Figure 1).

### Follow-up analyses

All follow up work focused on the targeted set of genes identified by the significant SGS regions. Genes were considered within significant segments for follow up if coding or regulatory sequence (defined using Genomic Regions Enrichment of Annotations Tool^21^) fell within the shared segment. Genes determined in previous research to be highly likely to represent false positive results^22^ were deleted (in our data, a cluster of 56 olfactory receptor genes in one region on chromosome 11, *FAT1, CTC-432M15.3*, and *TRIM51*). Three phases of follow-up work were pursued (see Figure S1).

### Corroborating evidence from the literature

As a first investigation of the genes indicated by significant SGS regions, we conducted a comprehensive search of the literature for all suicide-related risk genes against which to compare with SGS location-specific evidence. Suicide risk was identified by searching the Web of Science database for the terms: *suicid\** and *gene\** (captures variants including *suicide, suicidal, suicidality*, *gene, genetic,* etc.). We included reviews on the genetics of suicide as well as linkage, GWAS, candidate gene, expression and epigenetic studies. As secondary information about the suicide-related genes, we also queried DisGeNET^23,24^ for gene associations or involvement with neuropsychiatric disorders and inflammation due to their known association with suicide risk.^9,10,30^

### PsychArray variants in cases subsets in high-risk families with SGS sharing

The next phase of follow up comprised a search of the specific familial cases that generated each SGS region using available array variants within each region to determine if any particular array variant could be responsible each result. While the variants available to us on the PsychArray are far from complete, a search of relatively rare coding-region variants provides an efficient, immediate, potentially interpretable screen of our results in lieu of large-scale sequencing data.^26^ We therefore checked for sharing of the minor allele of non-synonymous variants within specific cases responsible for SGS results, and strictly within region boundaries. Because the SGS method is most powerful for the detection of rare familial variants, we selected a minor allele frequency <10% in the publicly-available Exome Aggregation Consortium (ExAC, www.exac.broadinstitute.org) European, non-Finnish data (matching in ancestry to our sample).

### Gene-based evidence in additional Utah suicide cases

The final follow-up phase focused on genes from SGS regions as targets for further study in additional sample resources. While familial variants may be private to the extended family/families producing the SGS evidence, it is also possible that the evidence implicates genes with additional risk variants in independent case samples. We screened an independent cohort of Utah suicides for potential functional variants in SGS-targeted genes; this case sample most closely matches the discovery families, as it was derived from the Utah population, and is comprised of completed suicides. Due to the same population ascertainment source, it is predicted to match the familial discovery sample regarding demographics and diagnostics. To maximize statistical power in our relatively small follow-up cohort of 1300 completed suicide cases, we focused on the potential to discover moderately penetrant, potentially interpretable functional causal variation. To this end, we used the following criteria to select variants: 1) in coding sequence of genes identified by the significant SGS regions, 2) nonsynonymous and predicted to be damaging from either PolyPhen^27^ or Sift,^28^ 3) minor allele frequency ≤ 20% in ExAC European, non-Finnish data. We tested for significant allelic association compared to ExAC European, non-Finnish data using Fisher’s exact test, or with chi-square tests for variants with >10 observed chromosomes with the minor allele in cases and controls. Tests for additional variants within SGS regions excluded suicide cases responsible for original sharing evidence in that region. Significance was adjusted for multiple tests.

## Results

### High-risk famlies

SGS analyses were performed on all 43 families. Most of the families (35/43=81.4%) showed at least one genome-wide suggestive region, and 10 families (23.3%) showed at least one genome-wide significant region. High-risk genealogies have additional complexity. Although the total number of cases across all families listed in Table 1 is 267, 52 of these cases occurred in multiple families (see Figure S3 for an example of this complexity). Because analyses to identify genomic shared regions are done within family, we included cases each time they occurred under each founder, as we don’t know *a priori* where true sharing may occur. It is possible that cases share risk variant(s) from one set of founders with other cases in that family, but then also share other risk variant(s) with cases in a second family through connections with the other founding couple. The complexities in relationships may allow for future studies of gene x gene interactions once risk variants have been established.

Descriptive characteristics of the 215 independent discovery cases from the 43 families were compared with the other 4,370 unselected Utah suicide cases with DNA. Within the 215 high-risk familial cases, 172 were male (80.0%), similar to the 79.2% rate in the unselected sample. Average age at death in the family sample was 34.28 years (standard deviation=16.28), significantly lower than the average age of 40.01 years (standard deviation=17.39) in the unselected sample (t=4.74; p<0.0001). Method of suicide in the family sample was predominantly gun-related (110/215=51.1%), followed by other violent methods (78/215=36.3%), then overdose (27/215=12.6%). These rates are similar to those in the unselected sample of 52.6%, 32.0%, and 15.3%, respectively. Death certificate data identified 212/215 cases as European-Non-Hispanic, and three as African-Non-Hispanic (one case each in families 41469, 233769, and 587072), similar to the unselected sample, where death certificate data identified 96.89% as European-Non-Hispanic. An ancestry principal component analysis of genotype data from the 215 cases and 1000 Genomes population data confirmed the three African ancestry cases (two showed Hispanic admixture), and identified 11 other cases with evidence of Asian and/or Hispanic ancestry, resulting in overall rates of 3.3% non-European and 5.6% Hispanic cases.

We found similar percentages of cases with presence of diagnoses from electronic records in the 215 familial cases as compared to the unselected 4,370 cases, with the exception of an increase in cases with personality disorders and an increase in prior attempts/suicidal ideation in the familial sample. Percentages of cases with at least one code in each of the diagnostic categories vs. the unselected sample were as follows: depression, 41.4% vs. 36.2%; bipolar, 13.0% vs. 10.6%; anxiety, 23.3% vs. 22.8%; psychosis, 2.3% vs. 2.9%; substance use/abuse, 9.8% vs. 12.4%; personality disorders, 14.4% vs. 9.7% (chi-square=5.06, p=0.02); ADHD, 4.7% vs. 3.7%; previous attempts/ideation, 37.2% vs. 28.9% (chi-square=6.82, p=0.009).

### Shared Genomic Segments (SGS) results

SGS analyses revealed 16 single-family regions with genome-wide significance (Table 2). Several families generated more than one region; these were the larger families, where there was more opportunity for multiple different case subsets to show sharing evidence (see Figure 1 for a specific example of sharing in family 66494; see Supplemental Figure S4a for drawings of all families with genome-wide significance). Table 2 also presents 15 regions where sharing evidence overlapped across more than one family (Figure S4b); in each of these regions, the single-family evidence was at least at the genome-wide suggestive level. For the region on chromosome 5q23.3-q31.1, person 112304 is a descendant of both 553615 and 603471, and person 95765 is a descendant in both 553615 and 176860. To satisfy the independence requirement for computing the Fisher’s combined p-value,^20^ we computed the p-value for this region omitting these cases. There are 207 genes with coding or regulatory sequence in the 31 SGS regions (Table S2).

### Follow up studies (see Figure S1 for overview)

#### 1) Supporting literature evidence

We did not find any overlap between significant SGS regions and genomic regions identified by previous family-based linkage studies of suicidal behaviors (Table S3). At the gene level, we reviewed the 207 SGS-targeted genes, first investigating specific supporting evidence of suicide risk. From our comprehensive literature search, a total of 755 genes have been associated with suicide with varying levels of statistical support (Table S4). Eighteen SGS-targeted genes were among these 755 suicide-risk genes (see Table 3; also highlighted in Table S2; a detailed description of these 18 genes follows Table S2). Given an estimated number of ~19,000 genes in the human genome,^29^ we estimate that 755/19000 = 4% of genes in the genome have current evidence associated with suicide risk. If the SGS regions were a random sample of the genome and unassociated with the suicide phenotype, we would expect that only about 4% of the genes in SGS regions (8/207 genes) would have corroborating evidence from the literature. However, we found that 18/207=8.7% of genes had supporting literature evidence, a significantly greater number than expected by chance (Z=2.41, p=0.008). This result suggests that the SGS regions are indeed segments of the genome that are enriched for prior evidence of suicide risk.

**Table 3.**
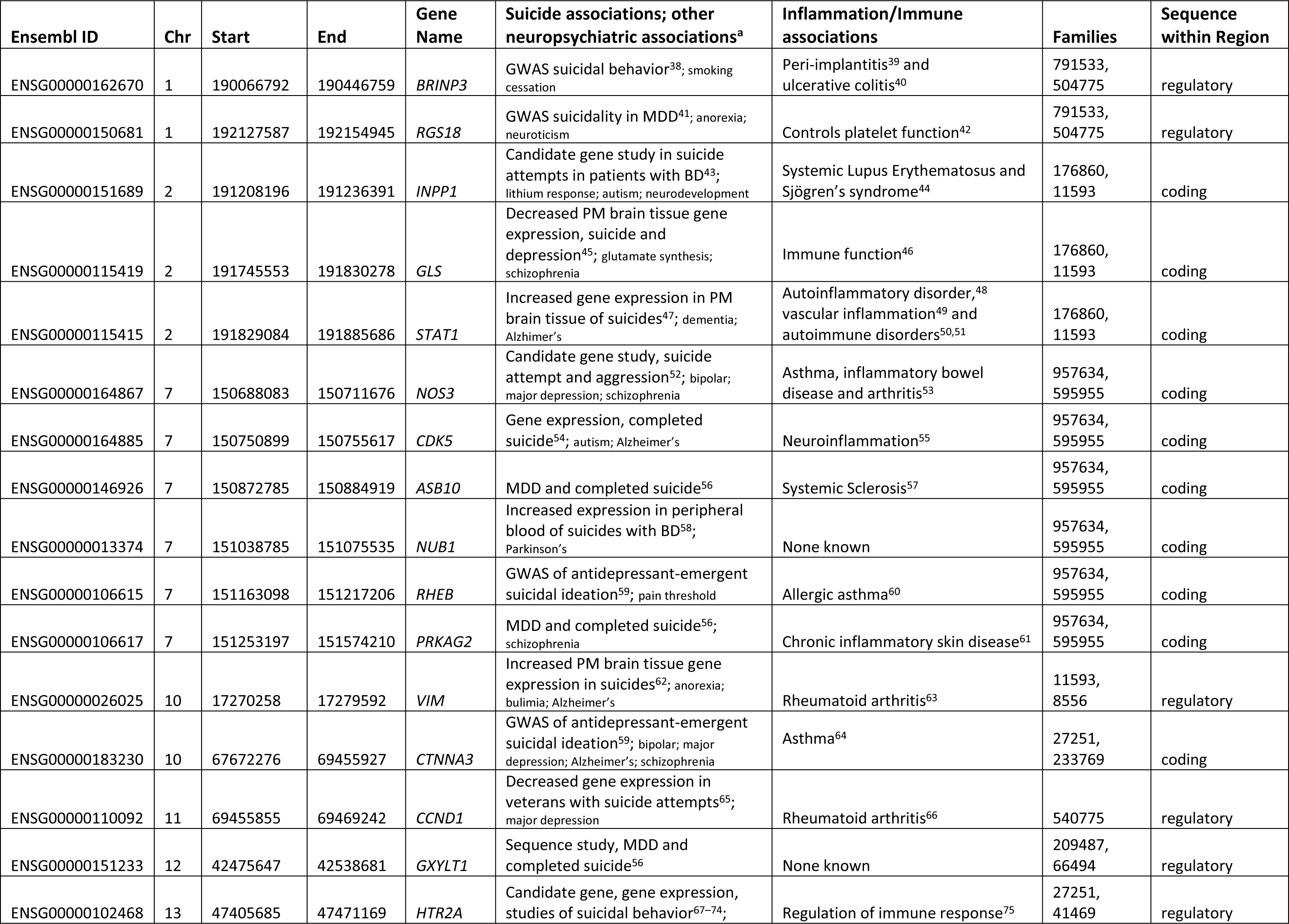
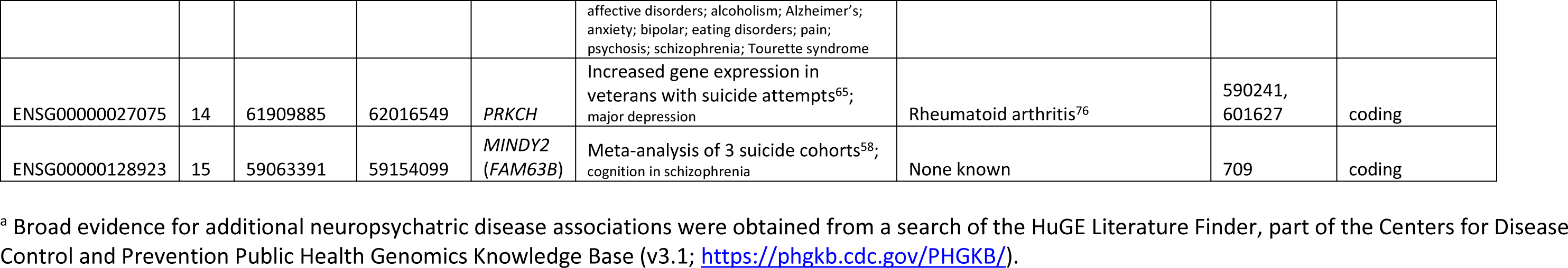
Genes in significant SGS regions with supporting evidence of association with suicide.

#### 2) Studies of variants in specific cases giving SGS evidence

We selected the 431 non-synonymous PsychArray variants with either benign or damaging functional predictions falling strictly within the significant SGS regions with ExAC European, non-Finnish minor allele frequency of ≤10%, reflecting the greater power of SGS to detect more rare risk variants. Considering each group of familial cases supporting each SGS signal, we screened the SNPs strictly within each region for sharing of the selected rare, non-synonymous array variants. There was no instance where this limited array content explained the identified SGS sharing.

#### 3) Studies of variants in ~1300 independent population-based cases

Additional population-ascertained Utah suicides with PsychArray genotype data were available for follow up. This sample was well-matched to the family discovery sample; a comparison of these two genotyped samples resulted in no significant demographic or diagnostic differences. These comparisons included the three variables that showed significant differences in our comparison between the family discovery sample and the larger Utah cohort of 4,370 suicides with DNA described above. Specifically, in the follow-up sample, age at death was 34.96 (standard deviation=16.76), percentage with personality disorders was 15.3%, and percentage with suicidal ideation/previous attempt was 36.9% (compared to 34.28 years, 14.4%, and 37.2%, respectively in the discovery family sample).

When we analyzed the selected 352 potentially damaging, relatively rare array variants within genes targeted by significant SGS regions, we found four variants with significantly increased presence of the minor allele compared to ExAC European non-Finnish frequencies, adjusting for multiple testing correction (Table 4: rs181058279, p=5.45E-06; rs76215382, p=8.48E-05; rs121908538, p=3.14E-12; rs745918508, p=5.40E-29). Specific characteristics of cases with each of these rare variants are described in Table S5. This evidence suggests rates of psychopathology similar to rates seen in the overall follow-up genotyped sample. Demographics were also similar, though cases with the *AGBL2* variant were significantly more likely to be female (chi-square=7.82, p=0.003).

**Table 4.** Putatively functional SNPs in target SGS genes with significantly elevated minor allele frequency in Utah suicide cases.

| Gene | SNP | Location<sup>a</sup> | change | Sift/Polyphen | ExAC Euro non-Finn freq (chroms)<sup>b</sup> | Suicide freq (chroms)<sup>c</sup> | p-value<sup>d</sup> | Function; disease association |
| --- | --- | --- | --- | --- | --- | --- | --- | --- |
| *SP110* | rs181058279 | chr2: 231033860 | G:C; missense | damaging; possibly damaging | 0.00006 (4/66714) | 0.0019 (5/2624) | 5.45E-06 | Gene transcription;<sup>30</sup> immune deficiency<sup>31,32</sup> |
| *AGBL2* | rs76215382 | chr11: 47711820 | G:A; missense | damaging; probably damaging | 0.0148 (986/66668) | 0.0247 (65/2622) | 8.48E-05 | ATP/GTP binding; brain structure and function<sup>33</sup> |
| *SUCLA2* | rs121908538 | chr13: 48528645 | A:G; missense | damaging; probably damaging | 0.00003 (2/66524) | 0.0034 (9/2621) | 3.14E-12 | Mitochondrial protein, energy to synapse<sup>34</sup> |
| *APH1B* | rs745918508 | chr15: 63594615 | -:A; frameshift | LOF | 0.00007 (5/66712) | 0.0088 (23/2620) | 5.40E-29 | Transmembrane protein; Alzheimer's<sup>36</sup> and Parkinson's<sup>37</sup> diseases |

<sup>a</sup> Base pair location hg19 genome build.
<sup>b</sup> Minor allele frequency and number of chromosomes with the minor allele from European non-Finnish samples in the Exome Aggregation Consortium (ExAC) data.
<sup>c</sup> Minor allele frequency and number of chromosomes with the minor allele derived from 1294-1312 Utah suicide cases with Illumina PsychArray data. For each variant, cases responsible for the original sharing were excluded. No case with a rare allele among those shown in this table had a known relationship (< 15<sup>th</sup> degree of relatedness) to the original high-risk family with SGS evidence.
<sup>d</sup> Comparison of UT suicide cases to the ExAC data; Fisher's exact test used for rs181058279, rs121908538, and rs745918508; chi-square test used for rs76215382. P-values exceed the significance threshold of 1.42E-04 correcting for 352 multiple tests.

## Discussion

We have ascertained and studied a unique resource of 43 extended families at high risk for suicide. The design uses the distantly related, high-risk cases to magnify genetic effects, enrich for genetic homogeneity, and minimize shared environmental effects. Families were identified from cases sampled from population-wide ascertainment, resulting in a study design independent of specific psychiatric diagnosis. Cases in the high-risk families were significantly younger at death, by 5.73 years on average, perhaps reflecting enhanced familial genetic risk over and above accumulated environmental risks that may play a greater role in suicide at later ages. The follow-up sample of ~1300 genotyped cases more closely matched this familial discovery sample. The mean age at death was similarly young, 5.05 years younger than the unelected sample. The matching of the replication cohort may be due to the fact that we have thus far targeted our overall genotyping efforts to cases with increased evidence of at least one other extended relative who is a suicide. Diagnostic data, when present, suggested similar rates of psychopathology across our entire research resource, with somewhat elevated rates of personality disorders and of suicidal ideation and previous attempt in both the family discovery sample and the follow-up genotyped sample.

Cases in families were analyzed with a statistically powerful method, SGS,^18^ resulting in the identification of genome-wide significant regions likely to harbor risk variants. This family evidence implicated 207 genes for targeted follow up. We found significant overlap with a comprehensive survey of 18 genes implicated in suicide, lending further support for these genes. Of note, 15 of these 18 genes also show previous associations with inflammatory conditions (Table 3), supporting accumulating evidence for a cross-association between inflammation and suicide risk.^25^ Because our method discovers familial genomic regions, we also reviewed prior family linkage studies of suicide risk, but did not find overlaps. This result is perhaps not surprising due to differences in ascertainment and outcome measures in these previous studies.

The additional rare disease-associated content of the array did not immediately reveal functional rare variants shared across cases responsible for the familial sharing. This result is likely due to the limited number of potentially risk-causing variants captured on the array; sequencing will be required to discover the causal variants shared across the high-risk discovery cases. Alternatively, one or more regions may be false positives.

SGS also provides target genes for other follow-up studies. Genes truly associated with suicide risk may harbor multiple risk-associated variants. By focusing our follow-up studies to the much reduced number of high-interest target variants in genes identified by SGS, statistical power is increased. An independent population-based cohort of ~1300 Utah completed suicide cases, well matched for ascertainment, resulted in four variants associated with suicide (*SP110*, *AGBL2*, *SUCLA2*, and *APH1B*). *SP110* is part of a leukocyte-specific nuclear body protein complex, and likely plays a role in gene transcription.^30^ It has been implicated in pathogen resistance and immunodeficiency,^31,32^ and may relate to suicide risk through a growing body of evidence implicating immune risk and inflammation.^25^ *AGBL2* is an ATP/GTP binding protein implicated in brain structure and function.^33^ S*UCLA2* is a mitochondrial tricarboxylic acid cycle protein recently implicated in energy supply to the synapse,^34^ and is possibly associated with recent findings linking suicide risk and hypoxia.^35^ *APH1B* is a transmembrane protein associated with risk of Alzheimer’s^36^ and Parkinson’s^37^ diseases. Characteristics of cases with each of these four variants did not reveal any striking patterns of association with specific psychopathology, though cases with the *AGBL2* variant were significantly more likely to be female. Follow up in additional research cohorts will be required to clarify diagnostic associations, and to replicate the association with gender found with the *AGBL2* variant.

### Limitations

Suicide cases were predominantly of Northern European ancestry, as verified with genotype data, so results may be limited to this race/ethnic group. The genome-wide background of the PsychArray contains ~265,000 common variants, which is relatively sparse for a genome-wide array. A denser array could have provided additional precision to region boundaries, or may have revealed that some regions were false positives. Diagnostic data were limited to available diagnoses in the electronic medical record. Cases with no diagnostic data are not assumed to have an absence of psychopathology. Rather, missing data more likely reflect either diagnoses outside the scope of our data resources, or lack of connection to services due to insurance, stigma, cultural factors or a perception that symptoms did not warrant treatment.

### Conclusions

While our study has found significant associations using only on the relatively rare, potentially functional variants captured on the PsychArray, these results have been discovered through a rigorous statistical prioritization and variant selection based only on functional annotation and frequency. As new data on our resource become available, it is likely that additional potential risk variation will be found. However, the current work has produced several important lines of evidence. First, the genome-wide significant SGS regions identify 207 target genes for suicide risk. Second, follow-up analyses of these regions in an independent population-based cohort of suicides highlighted four genes with potential functional risk variants, pending replication. Finally, the SGS regions contained 18 genes with corroborating evidence for suicide risk, suggesting these as strong candidates for future work.

## Acknowledgments

HC had full access to all the data in the study and takes responsibility for the integrity of the data and the accuracy of the data analysis. HC, LJ and ED are partially supported by a research contract from Janssen Research, LLC. QSL is an investigator at Janssen Research, LLC. No other authors have conflicts of interest relevant to the content of this manuscript, including no financial interest, relationships or affiliations. This work was supported by: the National Institute of Mental Health, R01MH099134 (HC) and K01MH093731 (AD), the American Foundation for Suicide Prevention (AVB), a Brain & Behavior Research Foundation Young Investigator Award (AD), the Clark Tanner Foundation (HC, TD, AVB), a donation from the Sharon Kae Lehr Endowed Research Fund in Memory of James Raymond Crump (DG), and the University of Utah EDGE Scholar Program (AD). Processing of samples was done with assistance from GCRC M01-RR025764 from the National Center for Research Resources. Partial support for all data sets within the UPDB was provided by the University of Utah Huntsman Cancer Institute. We especially thank University of Utah and OME staff whose hours of work made this study possible.

## Supporting Information Legends

**Table S1.** Diagnostic codes comprising possible co-occurring psychiatric conditions considered in this study. A case was defined as having evidence for a condition if one or more codes from the list below was present in the available electronic medical records data

**Table S2**. 207 genes with coding or regulatory sequence in regions with significance evidence for familial sharing from SGS analyses. Genes supporting evidence of suicide risk are highlighted in yellow. Psychiatric and/or neuronal associations are in bold type.

**Table S3**. Significant SGS regions in relation to previous linkage studies of suicidal ideation and behaviors.

**Table S4**. Genes with existing evidence for association with suicide risk from a comprehensive literature search.

**Table S5**. Case characteristics of follow-up Utah suicides with significant SNP findings. No cases were related (out to 15^th^ degree) to any case in the high risk family responsible for SGS evidence for the region containing that gene.

**Figure S1.** Flow diagram of the study.

**Figure S2**. Simplified hypothetical data defining familial shared genomic segments.

**Figure S3.** Example of suicide cases with complex family relationships to more than one founding couple.

**Figure S4a**. Extended family structures linking cases used for SGS analyses in the 10 families with genome-wide significant results within a single family. All analyzed cases are represented by a numeric ID. Only suicide cases in the line of descent to analyzed cases are shown. Gender is disguised and sibship order is randomized in order to protect the privacy of family members. NOTE: Suicide cases are not as evident in upper generations because suicide status from death certificates is only available back to 1904.

**Figure S4b**. Detail of extended family structures in the families with SGS evidence of shared regions overlapping across more than one family. All regions shared across two families are presented in drawings i through xiv. The region on chromosome five, shared across three families, is presented in drawing xv.

## References

1 Centers for Disease Control & Prevention (CDC) Data & Statistics Fatal Injury Report for 2015. 2015. https://webappa.cdc.gov/sasweb/ncipc/mortrate.html.

2 Pedersen NL, Fiske A. Genetic influences on suicide and nonfatal suicidal behavior: Twin study findings. Eur Psychiatry 2010; 25: 264–267.

3 McGuffin P, Marusic A, Farmer A. What can psychiatric genetics offer suicidology? Crisis. 2001; 22: 61–65.

4 Mirkovic B, Laurent C, Podlipski M-A, Frebourg T, Cohen D, Gerardin P. Genetic Association Studies of Suicidal Behavior: A Review of the Past 10Years, Progress, Limitations, and Future Directions. Front Psychiatry 2016; 7. doi: 10.3389/fpsyt.2016.00158.

5 Younes N, Melchior M, Turbelin C, Blanchon T, Hanslik T, Chan Chee C. Attempted and completed suicide in primary care: Not what we expected? J Affect Disord 2015; 170: 150–154.

6 Groden J, Thliveris a, Samowitz W, Carlson M, Gelbert L, Albertsen H et al. Identification and characterization of the familial adenomatous polyposis coli gene. Cell 1991; 66: 589–600.

7 Goldgar DE, Cannon-Albright L a, Oliphant a, Ward JH, Linker G, Swensen J et al. Chromosome 17q linkage studies of 18 Utah breast cancer kindreds. Am J Hum Genet 1993; 52: 743–8.

8 Curran ME, Splawski I, Timothy KW, Vincent GM, Green ED, Keating MT. A molecular basis for cardiac arrhythmia: HERG mutations cause long QT syndrome. Cell 1995; 80: 795–803.

9 Arsenault-Lapierre G, Kim C, Turecki G. Psychiatric diagnoses in 3275) suicides: A meta-analysis. BMC Psychiatry. 2004; 4. doi:10.1186/1471-244X-4-37.

10 Brezo J, Paris J, Tremblay R, Vitaro F, Hébert M, Turecki G. Identifying correlates of suicide attempts in suicidal ideators: A population-based study. Psychol Med 2007; 37: 1551–1562.

11 McGirr A, Alda M, Séguin M, Cabot S, Lesage A, Turecki G. Familial aggregation of suicide explained by cluster B traits: A three-group family study of suicide controlling for major depressive disorder. Am J Psychiatry 2009; 166: 1124–1134.

12 Kim CD, Seguin M, Therrien N, Riopel G, Chawky N, Lesage AD et al. Familial aggregation of suicidal behavior: A family study of male suicide completers from the general population. Am J Psychiatry 2005; 162: 1017–1019.

13 Brent DA, Oquendo M, Birmaher B, Greenhill L, Kolko D, Stanley B et al. Familial pathways to early-onset suicide attempt: risk for suicidal behavior in offspring of mood-disordered suicide attempters. Arch Gen Psychiatry 2002; 59: 801–7.

14 Egeland JA, Sussex JN. Suicide and Family Loading for Affective Disorders. JAMA J Am Med Assoc 1985; 254: 915–918.

15 Geschwind DH, Flint J. Genetics and genomics of psychiatric disease. Science 2015; 349: 1489–1494.

16 Kerber RA. Method for calculating risk associated with family history of a disease. Genet Epidemiol 1995; 12: 291–301.

17 Chang CC, Chow CC, Tellier LCAM, Vattikuti S, Purcell SM, Lee JJ. Second-generation PLINK: Rising to the challenge of larger and richer datasets. Gigascience 2015; 4. doi:10.1186/s13742-015-0047-8.

18 Waller RG, Darlington TM, Wei X, Madsen MJ, Thomas A, Curtin K et al. Novel pedigree analysis implicates DNA repair and chromatin remodeling in multiple myeloma risk. PLoS Genet 2018; 14. doi:10.1371/journal.pgen.1007111.

19 Knight S, Abo RP, Abel HJ, Neklason DW, Tuohy TM, Burt RW et al. Shared Genomic Segment Analysis: The Power to Find Rare Disease Variants. Ann Hum Genet 2012; 76: 500–509.

20 Fisher R. Statistical methods for research workers. 1925 doi:10.1056/NEJMc061160.

21 McLean CY, Bristor D, Hiller M, Clarke SL, Schaar BT, Lowe CB et al. GREAT improves functional interpretation of cis-regulatory regions. Nat Biotechnol 2010; 28: 495–501.

22 Fuentes Fajardo K V., Adams D, Mason CE, Sincan M, Tifft C, Toro C et al. Detecting false-positive signals in exome sequencing. Hum Mutat 2012; 33: 609–613.

23 Piñero J, Bravo Á, Queralt-Rosinach N, Gutiérrez-Sacristán A, Deu-Pons J, Centeno E et al. DisGeNET: A comprehensive platform integrating information on human disease-associated genes and variants. Nucleic Acids Res 2017; 45: D833–D839.

24 Piñero J, Queralt-Rosinach N, Bravo À, Deu-Pons J, Bauer-Mehren A, Baron M et al. DisGeNET: A discovery platform for the dynamical exploration of human diseases and their genes. Database 2015; 2015. doi:10.1093/database/bav028.

25 Brundin L, Bryleva EY, Thirtamara Rajamani K. Role of Inflammation in Suicide: From Mechanisms to Treatment. Neuropsychopharmacology. 2017; 42: 271–283.

26 Boyle EA, Li YI, Pritchard JK. An Expanded View of Complex Traits: From Polygenic to Omnigenic. Cell. 2017; 169: 1177–1186.

27 Adzhubei IA, Schmidt S, Peshkin L, Ramensky VE, Gerasimova A, Bork P et al. A method and server for predicting damaging missense mutations. Nat. Methods. 2010; 7: 248–249.

28 Kumar P, Henikoff S, Ng PC. Predicting the effects of coding non-synonymous variants on protein function using the SIFT algorithm. Nat Protoc 2009; 4: 1073–1082.

29 Ezkurdia I, Juan D, Rodriguez JM, Frankish A, Diekhans M, Harrow J et al. Multiple evidence strands suggest that theremay be as few as 19 000 human protein-coding genes. Hum Mol Genet 2014; 23: 5866–5878.

30 Leu J, Chang S, Mu C, Chen M, Yan B. Functional domains of SP110 that modulate its transcriptional regulatory function and cellular translocation. J Biomed Sci 2018; 25: 34.

31 Marquardsen FA, Baldin F, Wunderer F, Al-Herz W, Mikhael R, Lefranc G et al. Detection of Sp110 by Flow Cytometry and Application to Screening Patients for Veno-occlusive Disease with Immunodeficiency. J Clin Immunol 2017; 37: 707–714.

32 Cai L, Wang Y, Wang J-F, Chou K-C. Identification of Proteins Interacting with Human SP110 During the Process of Viral Infections. Med Chem (Los Angeles) 2011; 7: 121–126.

33 Karaca E, Harel T, Pehlivan D, Jhangiani SN, Gambin T, Coban Akdemir Z et al. Genes that Affect Brain Structure and Function Identified by Rare Variant Analyses of Mendelian Neurologic Disease. Neuron 2015; 88: 499–513.

34 Völgyi K, Gulyássy P, Háden K, Kis V, Badics K, Kékesi KA et al. Synaptic mitochondria: A brain mitochondria cluster with a specific proteome. J Proteomics 2015; 120: 142–157.

35 Kious BM, Kondo DG, Renshaw PF. Living High and Feeling Low: Altitude, Suicide, and Depression. Harv Rev Psychiatry 2018; 26: 43–56.

36 Haghighi F, Xin Y, Chanrion B, O’Donnell AH, Ge Y, Dwork AJ et al. Increased DNA methylation in the suicide brain. Dialogues Clin Neurosci 2014; 16: 430–438.

37 Bekris LM, Tsuang DW, Peskind ER, Yu CE, Montine TJ, Zhang J et al. Cerebrospinal fluid Abeta42 levels and APP processing pathway genes in Parkinson’s disease. Mov Disord 2015; 30: 936–944.

38 Galfalvy H, Haghighi F, Hodgkinson C, Goldman D, Oquendo MA, Burke A et al. A genome-wide association study of suicidal behavior. Am J Med Genet Part B Neuropsychiatr Genet 2015; 168: 557–563.

39 Casado PL, Aguiar DP, Costa LC, Fonseca MA, Vieira TCS, Alvim-Pereira CCK et al. Different contribution of BRINP3 gene in chronic periodontitis and peri-implantitis: A cross-sectional study. BMC Oral Health 2015; 15. doi:10.1186/s12903-015-0018-6.

40 Smith PJ, Levine AP, Dunne J, Guilhamon P, Turmaine M, Sewell GW et al. Mucosal transcriptomics implicates under expression of BRINP3 in the pathogenesis of ulcerative colitis. Inflamm Bowel Dis 2014; 20: 1802–1812.

41 Schosser A, Butler AW, Ising M, Perroud N, Uher R, Ng MY et al. Genomewide association scan of suicidal thoughts and behaviour in major depression. PLoS One 2011; 6. doi:10.1371/journal.pone.0020690.

42 Xie Z, Chan EC, Druey KM. R4 Regulator of G Protein Signaling (RGS) Proteins in Inflammation and Immunity. AAPS J 2016; 18: 294–304.

43 Jiménez E, Arias B, Mitjans M, Goikolea JM, Roda E, Sáiz PA et al. Genetic variability at IMPA2, INPP1 and GSK3β increases the risk of suicidal behavior in bipolar patients. Eur Neuropsychopharmacol 2013; 23: 1452–1462.

44 Lindén M, Ramírez Sepúlveda JI, James T, Thorlacius GE, Brauner S, Gómez-Cabrero D et al. Sex influences eQTL effects of SLE and Sjögren’s syndrome-associated genetic polymorphisms. Biol Sex Differ 2017; 8. doi:10.1186/s13293-017-0153-7.

45 Fiori LM, Turecki G. Broadening our horizons: Gene expression profiling to help better understand the neurobiology of suicide and depression. Neurobiol. Dis. 2012; 45: 14–22.

46 Xu FF, Huang Y, Wang XQ, Qiu YH, Peng YP. Modulation of immune function by glutamatergic neurons in the cerebellar interposed nucleus via hypothalamic and sympathetic pathways. Brain Behav Immun 2014; 38: 263–271.

47 Hoyo-Becerra C, Huebener A, Trippler M, Lutterbeck M, Liu ZJ, Truebner K et al. Concomitant interferon alpha stimulation and TLR3 activation induces neuronal expression of depression-related genes that are elevated in the brain of suicidal persons. PLoS One 2013; 8. doi:10.1371/journal.pone.0083149.

48 Liu Y, Ramot Y, Torrelo A, Paller AS, Si N, Babay S et al. Mutations in proteasome subunit β type 8 cause chronic atypical neutrophilic dermatosis with lipodystrophy and elevated temperature with evidence of genetic and phenotypic heterogeneity. Arthritis Rheum 2012; 64: 895–907.

49 Chmielewski S, Piaszyk-Borychowska A, Wesoly J, Bluyssen HAR. STAT1 and IRF8 in Vascular Inflammation and Cardiovascular Disease: Diagnostic and Therapeutic Potential. Int Rev Immunol 2016; 35: 434–454.

50 Chen Z, Guo Z, Ma J, Liu F, Gao C, Liu S et al. STAT1 single nucleotide polymorphisms and susceptibility to immune thrombocytopenia. Autoimmunity 2015; 48: 305–312.

51 Zimmerman O, Rosen LB, Swamydas M, Ferre EMN, Natarajan M, van de Veerdonk F et al. Autoimmune regulator deficiency results in a decrease in STAT1 levels in human monocytes. Front Immunol 2017; 8. doi:10.3389/fimmu.2017.00820.

52 Giegling I, Calati R, Porcelli S, Hartmann AM, Möller HJ, De Ronchi D et al. NCAM1, TACR1 and NOS genes and temperament: A study on suicide attempters and controls. Neuropsychobiology 2011; 64: 32–37.

53 Kanwar JR, Kanwar RK, Burrow H, Baratchi S. Recent advances on the roles of NO in cancer and chronic inflammatory disorders. Curr Med Chem 2009; 16: 2373–94.

54 Choi K, Le T, Xing G, Johnson LR, Ursano RJ. Analysis of Kinase Gene Expression in the Frontal Cortex of Suicide Victims: Implications of Fear and Stress†. Front Behav Neurosci 2011; 5. doi:10.3389/fnbeh.2011.00046.

55 Quintanilla RA, Orellana DI, González-Billault C, Maccioni RB. Interleukin-6 induces Alzheimer-type phosphorylation of tau protein by deregulating the cdk5/p35 pathway. Exp Cell Res 2004; 295: 245–257.

56 Tombácz D, Maróti Z, Kalmár T, Csabai Z, Balázs Z, Takahashi S et al. High-Coverage Whole-Exome Sequencing Identifies Candidate Genes for Suicide in Victims with Major Depressive Disorder. Sci Rep 2017; 7: 1–11.

57 Gao L, Emond MJ, Louie T, Cheadle C, Berger AE, Rafaels N et al. Identification of Rare Variants in ATP8B4 as a Risk Factor for Systemic Sclerosis by Whole-Exome Sequencing. Arthritis Rheumatol 2016; 68: 191–200.

58 Niculescu AB, Le-Niculescu H, Levey DF, Phalen PL, Dainton HL, Roseberry K et al. Precision medicine for suicidality: From universality to subtypes and personalization. Mol Psychiatry 2017; 22: 1250–1273.

59 Menke A, Domschke K, Czamara D, Klengel T, Hennings J, Lucae S et al. Genome-wide association study of antidepressant treatment-emergent suicidal ideation. Neuropsychopharmacology 2012; 37: 797–807.

60 Li K, Zhang Y, Liang KY, Xu S, Zhou XJ, Tan K et al. Rheb1 deletion in myeloid cells aggravates OVA-induced allergic inflammation in mice. Sci Rep 2017; 7. doi:10.1038/srep42655.

61 Shen C, Liu L, Jiang Z, Zheng X, Meng L, Yin X et al. Four genetic variants interact to confer susceptibility to atopic dermatitis in Chinese Han population. Mol Genet Genomics 2015; 290: 1493–8.

62 Kékesi KA, Juhász G, Simor A, Gulyássy P, Szego ÉM, Hunyadi-Gulyás É et al. Altered Functional Protein Networks in the Prefrontal Cortex and Amygdala of Victims of Suicide. PLoS One 2012; 7. doi:10.1371/journal.pone.0050532.

63 Wegner N, Lundberg K, Kinloch A, Fisher B, Malmström V, Feldmann M et al. Autoimmunity to specific citrullinated proteins gives the first clues to the etiology of rheumatoid arthritis. Immunol. Rev. 2010; 233: 34–54.

64 Perin P, Potočnik U. Polymorphisms in recent GWA identified asthma genes CA10, SGK493, and CTNNA3 are associated with disease severity and treatment response in childhood asthma. Immunogenetics 2014; 66: 143–151.

65 Flory JD, Donohue D, Muhie S, Yang R, Miller SA, Hammamieh R et al. Gene expression associated with suicide attempts in US veterans. Transl Psychiatry 2017; 7. doi: 10.1038/tp.2017.179.

66 Katoh M, Katoh M. STAT3-induced WNT5A signaling loop in embryonic stem cells, adult normal tissues, chronic persistent inflammation, rheumatoid arthritis and cancer (Review). Int. J. Mol. Med. 2007; 19: 273–278.

67 Bani-Fatemi A, Howe AS, Matmari M, Koga A, Zai C, Strauss J et al. Interaction between methylation and CpG single-nucleotide polymorphisms in the HTR2A gene: Association analysis with suicide attempt in schizophrenia. Neuropsychobiology 2016; 73: 10–15.

68 González-Castro TB, Tovilla-Zárate C, Juárez-Rojop I, Pool García S, Velázquez-Sánchez MP, Genis A et al. Association of the 5HTR2A gene with suicidal behavior: CASE-control study and updated meta-analysis. BMC Psychiatry 2013; 13. doi: 10.1186/1471-244X-13-25.

69 Garbett K, Gal-Chis R, Gaszner G, Lewis DA, Mirnics K. Transcriptome alterations in the prefrontal cortex of subjects with schizophrenia who committed suicide. Neuropsychopharmacol Hung 2008; 10: 9–14.

70 Sequeira A, Morgan L, Walsh DM, Cartagena PM, Choudary P, Li J et al. Gene expression changes in the prefrontal cortex, anterior cingulate cortex and nucleus accumbens of mood disorders subjects that committed suicide. PLoS One 2012; 7. doi:10.1371/journal.pone.0035367.

71 Mann JJ, Stanley M, McBride PA, McEwen BS. Increased Serotonin2 and β-Adrenergic Receptor Binding in the Frontal Cortices of Suicide Victims. Arch Gen Psychiatry 1986; 43: 954–959.

72 Antypa N, Serretti A, Rujescu D. Serotonergic genes and suicide: A systematic review. Eur. Neuropsychopharmacol. 2013; 23: 1125–1142.

73 Mandelli L, Serretti A. Gene environment interaction studies in depression and suicidal behavior: An update. Neurosci. Biobehav. Rev. 2013; 37: 2375–2397.

74 Schild AHE, Pietschnig J, Tran US, Voracek M. Genetic association studies between SNPs and suicidal behavior: A meta-analytical field synopsis. Prog Neuro-Psychopharmacology Biol Psychiatry 2013; 46: 36–42.

75 Idzko M, Panther E, Stratz C, Müller T, Bayer H, Zissel G et al. The serotoninergic receptors of human dendritic cells: identification and coupling to cytokine release. J Immunol 2004; 172: 6011–9.

76 Takata Y, Hamada D, Miyatake K, Nakano S, Shinomiya F, Scafe CR et al. Genetic association between the PRKCH gene encoding protein kinase Ceta isozyme and rheumatoid arthritis in the Japanese population. Arthritis Rheum 2007; 56: 30–42.

